# The utility of visual estimation of cover for rapid assessment of graminoid abundance in forest and grassland habitats in studies of animal foraging

**DOI:** 10.1101/012716

**Authors:** Hansraj Gautam, G. G. Potdar, T.N.C. Vidya

## Abstract

**Questions:** To assess the feasibility of using visually-estimated cover in rapid assessment of herbivore food species abundance in the grass layer, we asked the following questions: 1) What is the relationship between total graminoid cover and biomass in forests, and does height improve the prediction of biomass from cover? 2) How does total cover relate to biomass in a grassland habitat? 3) How does elephant food species graminoid cover relate to individual species biomass? 4) How well does species diversity of forest understorey grass layer, calculated from cover data, mirror that calculated from biomass data?

**Location:** Nagarahole National Park, India.

**Methods:** We estimated the abundance of graminoids through visual estimation of cover and weighted harvested biomass in forest and grassland plots. In forests, two estimates of total graminoid abundance, total graminoid cover and sum of species covers, were used. In the grassland, only total graminoid abundance was measured. We examined the relationship between cover estimates and biomass, and the additional utility of height in predicting biomass, using multiple regressions and AIC-based model selection. We also assessed similarity in cover- and biomass-based Simpson’s and Shannon-Wiener diversity indices using regressions.

**Results:** Graminoid cover explained a large portion of variation in total graminoid biomass in forest and grassland habitats. The sum of species covers was better than total cover in estimating total graminoid biomass in the forest. The benefit of including height to estimate total biomass was moderate in forests but substantial in grassland. Cover estimates were good proxies of food species biomass, and the addition of height did not yield better models for most species. Species diversity indices calculated from cover largely matched those based on biomass.

**Conclusions:** Visual estimation of species cover is a good alternative to biomass harvesting for rapid assessment of abundance of graminoids consumed by generalist herbivores, like elephants.

## Introduction

Ecologists estimate vegetation abundance in order to study various structural and functional attributes of plant communities (e.g. Hermy 1988, Guo and Rundel 1997, Chiarucci et al. 1999, Henschel et al. 2005, Lavorel et al. 2008), the productivity of animals’ habitats (e.g. Hutto 1990, Säid et al. 2005, Pettorelli et al. 2011, Iversion et al. 2014) and its effect on foraging behaviour (e.g. Wilmshurst et al. 1999), and the impact of animal activities on vegetation (e.g. Pekin et al. 2015). While studies of plant community structure and function may require intensive measurements of species abundance or traits (e.g. Chiarucci et al. 1999, Lavorel et al. 2008), assessment of resource availability for animals often necessitates sampling over large spatial scales (see Pettorelli et al. 2011), which would, therefore, benefit from rapid methods of estimating species abundance. The estimation of forage abundance is a pre-requisite in studies of ecological and behavioural aspects of foraging ecology (Hutto 1990, Säid et al. 2005), but the collection of detailed forage abundance data may be very demanding in terms of effort, time, and resources, which are limitations for most field biologists. Different methods of estimating vegetation abundance vary in their sensitivity to vegetation structure, accuracy, precision, practicality, time and manpower required, and destructive nature (Harmoney et al. 1997, reviewed in Wilson 2011), and no single method is clearly superior in all these respects. Given the apparent trade-off in methods between accuracy and speed, it has been suggested that the choice of method should be based on the objective of the study and after consideration of the advantages and limitations of each method (Elzinga et al. 1998, Lavorel et al. 2008, Wilson 2011, Redjadj et al. 2012).

The diet of the animal and the vegetation structure in its habitat will together determine whether the measurement of forage abundance should be carried out on all components of the vegetation or only on a portion of the vegetation (such as specific plants or plant parts). For example, all the vegetation of a largely monocultural grassland may be considered for quantifying the food abundance of a grazing herbivore, whereas many components of vegetation (herbs, shrubs, and tree species) in a forest or woodland may not be part of the same animal’s diet, and only the food component of the vegetation should be measured. Second, in studies of foraging, ecologists are also often interested in studying whether an animal shows selectivity at the species level during feeding (e.g. Owen-Smith and Chafota 2012) and whether it maintains species diversity in its diet (e.g. Marsh et al. 2006). It is, therefore, also important for the method of estimation of forage abundance to provide species-level detail. Given the considerations above, several methods of abundance estimation become impractical or too time-consuming to implement in diverse habitats, primarily because the vegetation layers that are not relevant to foraging may dominate the biomass in forests with rich biodiversity. Unlike other estimation methods, the biomass harvest method (e.g. Drew 1944, Hermy 1988), and the visual estimation of cover (Kennedy and Addison 1987) can be applied even when a selected portion of vegetation is to be quantified. However, biomass-harvesting can be time-consuming if species have to be weighed separately, as this requires sorting of individuals into different species by hand (Harmony et al. 1997, Lavorel et al. 2008). In this regard, the use of visual estimation of cover may be advantageous, as it allows for rapid assessment of portions of vegetation and is also non-destructive.

We, therefore, tested the utility of the visual estimation method in predicting biomass in the context of forage availability for herbivores in general and for Asian elephants in a tropical forest in southern India in particular. Elephants are considered generalist herbivores, but primarily feed on grasses in the lower vegetation strata, in addition to stems and bark of woody species in the upper strata (Owen-Smith 1988, Sukumar 1990, Baskaran et al. 2010). Although their diet consists of numerous species, in species-rich tropical forests, this number may be a small proportion (Blake 2002, HG and TNCV unpubl. data) of all the species present. The estimation of abundance of woody species is simple as it involves the counting of trees and measurements of tree-girth which can be done rapidly since the number per plot is usually low. However, elephant foods in understorey vegetation are represented by numerous individuals that are difficult to count within limited time. Moreover, most of the vegetation represented by herbs and shrubs is not consumed by elephants (in our study area, the abundance of food species as a percentage of all species in the respective vegetation strata during the wet and dry seasons, respectively, was about 23% and 10% for herbs, 18% and 16% for shrubs and 80% and 85% for graminoids, HG and TNCV unpubl. data). Therefore, in such habitats, the focus should be on estimating the abundance of only food plant species in the lower strata of the forest. The dominance of grasses in Asian elephant diet (Baskaran et al. 2010) makes its quantification crucial, and we explored the utility of the visual estimation method in assessing graminoid biomass. Since other herbivores in similar deciduous forests are also primarily grazers (Ahrestani et al. 2012), if the visual method could be used to reliably estimate biomass, our results would also have implications for the quantification of resource abundance for such herbivores. Therefore, we assessed the utility of visually estimated cover in explaining elephant food graminoid biomass, as well as total graminoid biomass, which would establish the generality of the method, for use in other species. We investigated the utility of this rapid method at the community level, as well as at the more detailed species level. We also examined the additional utility of height, another variable which can be rapidly measured, in modelling graminoid biomass in two types of habitats, forest and grassland. Such questions regarding biomass of graminoids have been rarely addressed in forest habitats (eg. Andariese and Covington 1986), especially in the context of forage availability for wildlife.

Previously, studies have found strong correlations between visual estimates and biomass (Hermy 1988, Guo and Rundel 1997, Chiarucci et al. 1999, Henschel et al. 2005, Axmanová et al. 2012) but were not carried out in the context of sampling food availability for wildlife. On the other hand, studies on foraging ecology have sometimes used visual estimation for assessment of forage distribution (e.g. Noyce and Coy 1990, Blake 2002, Rebollo et al. 2013, Iversion et al. 2014), but the relationship between visual estimates and biomass of relevant foods has seldom been tested rigorously in a complex habitat (but see Noyce and Coy 1990 for bear foods), which is important before making inferences about the relationship between resource distribution and forage selection.

The questions we addressed in this paper were the following:

1. What is the relationship between visually-estimated total graminoid cover and total graminoid biomass (measured through the standard biomass-harvest method) in forest habitats, and does the inclusion of height or using the visually-estimated sum of species covers improve the prediction of total graminoid biomass? This question would help find out if the visual estimation method can be used in general in a forest with multiple strata, in studies of foraging by grazing herbivores.
2. Does the relationship between visually-estimated total graminoid cover and total graminoid biomass (as seen from the results of question 1) also hold in a grassland habitat, and does the inclusion of height improve the prediction of total graminoid abundance?
3. How do visually-estimated species covers of individual graminoid food species of elephants relate to their respective species biomass measurements in forest habitats? Since the proportion of food graminoid species represents only a small fraction of all species in the herbaceous stratum of the vegetation in the forest sampled (HG and TNCV unpubl. data), if there was a high correlation between visually-estimated species cover and species biomass, visually estimated cover could be used to assess food species abundance and also estimate proportional abundance of different species.
4. How accurately do species diversity indices of the grass layer in forest habitat, measured by visual estimates, reflect the diversity indices obtained from biomass data? It would be desirable to obtain good diversity estimates in order to study selectivity of species and selection of different kinds of vegetation patches by herbivores.

## Study area

The study was carried out in Nagarahole National Park (644 km^2^, 11.85°–12.26° N, 76.00°–76.28° E), which is a part of the larger contiguous Nilgiris-Eastern Ghats landscape in southern India (Figure 2). The forest is tropical deciduous, comprising several strata, and is home to several herbivores, including Asian elephants, on which a long-term study based on individually identified elephants is currently ongoing (see Vidya et al. 2014). Along the southern boundary of the park flows the river Kabini, on which a dam was constructed during the 1970s to create a reservoir that extends along the southern boundary of the park. During the dry season, when the waters of the reservoir recede, the exposed area forms a grassland consisting mostly of just two short grass species (*Cynodon dactylon* and *Sporobolus* sp., which are also found in the forest, see Supporting Information 1) and attracts a large number of elephants, deer, and gaur. The graminoids in the grassland are shorter (mean height 5.7 cm) and more continuously distributed compared to those in the forests (mean weighted average of species heights 24.6 cm), where they are more sparsely distributed. Both the grassland and the forest habitat are used by elephants, and data on graminoid abundance were collected from both types of habitats, as described below.

## Data sampling and analysis

### Forest data

Data collection in forests was done from November to December 2013. Based on a forest type classification map of the region developed by Pascal (1982), Nagarahole National Park was divided into the three major forest types: dry deciduous forest, moist deciduous forest, and teak plantations. We had previously divided the area into a 2 km × 2 km grid and placed 60 1-km line transects in randomly selected cells in order to map the distribution of elephant food resources. During the present study, 23 of these transects in the southern and central parts of the park and 17 additional transects, at least half a km away from each other and at least 100 m away from forest roads, were chosen for sampling. Care was taken to adequately represent all three forest types (based on their availability) in the sampling sites. Sampled locations are mapped in Figure 2.

Sampling was carried out in 20 m × 5 m plots at the start or end of each of the 40 1-km transects. In each of these 20 m × 5 m plots, three 1 m × 1 m quadrats were sampled, equidistant along a straight diagonal line (except one plot, in which only two quadrats could be sampled due to the presence of dense *Lantana* thickets). This resulted in 119 quadrats sampled. We sampled all graminoid plants, including *Poaceae* (grasses), *Juncaceae* and *Cyperaceae*. Graminoid abundance was measured at two levels: total graminoid abundance (all the graminoid species present) and species-level abundance. First, total graminoid cover was visually estimated by a single observer (HG) as the percentage of quadrat area covered by all graminoids (Figure 1). Second, species cover for each graminoid species was visually estimated, independent of the cover of other species. Cover was estimated to the closest 5% or in interval bin of 5% (for low values such as 0 to 5% cover), in which case, the middle value of the interval was chosen as the cover value. Values of less than 5% were applied in the case of rare species that were represented by only one or two individuals in the quadrat. Four individuals (except in the case of rare species, in which fewer than four individuals were available) of each species were arbitrarily selected, their natural standing heights (i.e. without straightening the plant) were measured, and the average of these was used as the height for that species. The total graminoid (fresh) biomass was measured in the field using a digital weighing balance (with 1 gram precision) after harvesting all the graminoids from the ground level. Individuals were then hand-sorted into the respective species, and the biomass of each species was measured.

**Figure 1:**
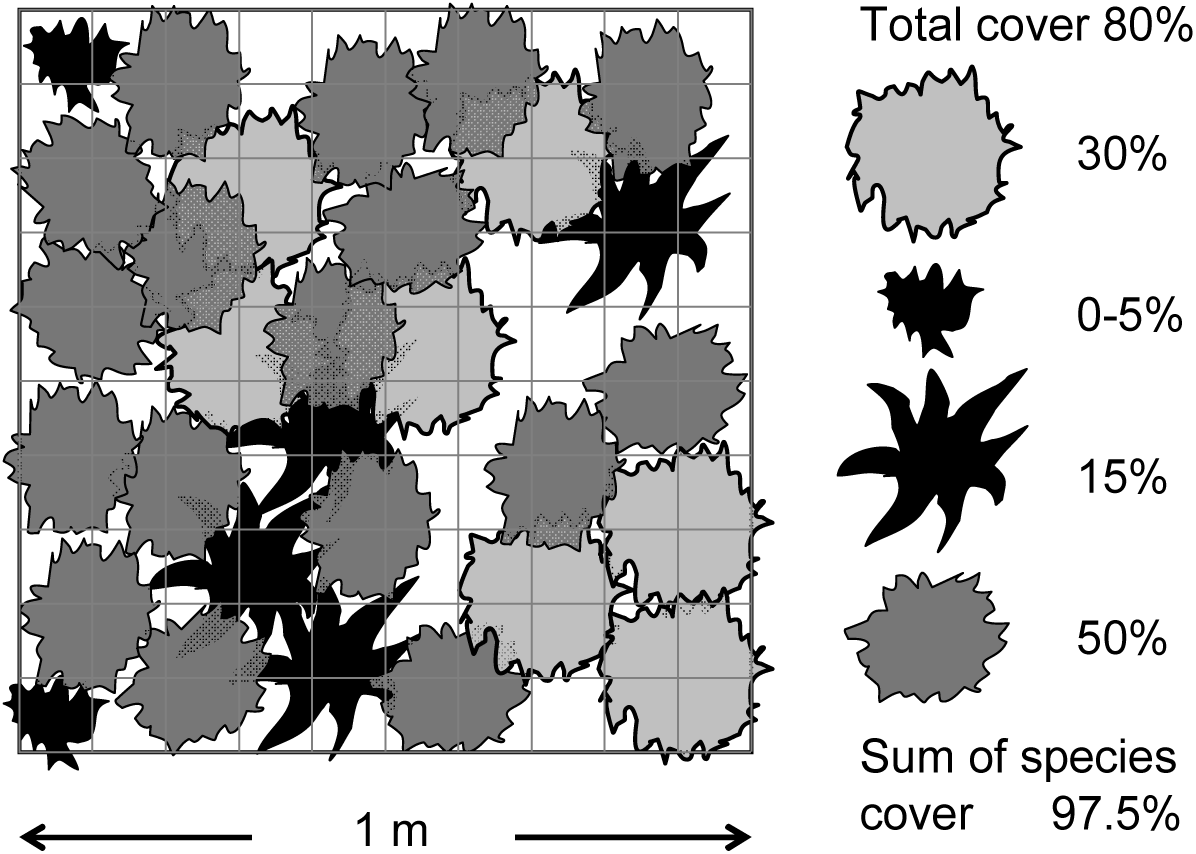
Illustration of a 1 m × 1 m sampling quadrat, showing estimates of total cover, individual species covers of four species (shown as different combinations of shape and colour), and sum of species cover. The sum of species cover is higher than total cover primarily because of between-species overlaps.

**Figure 2:**
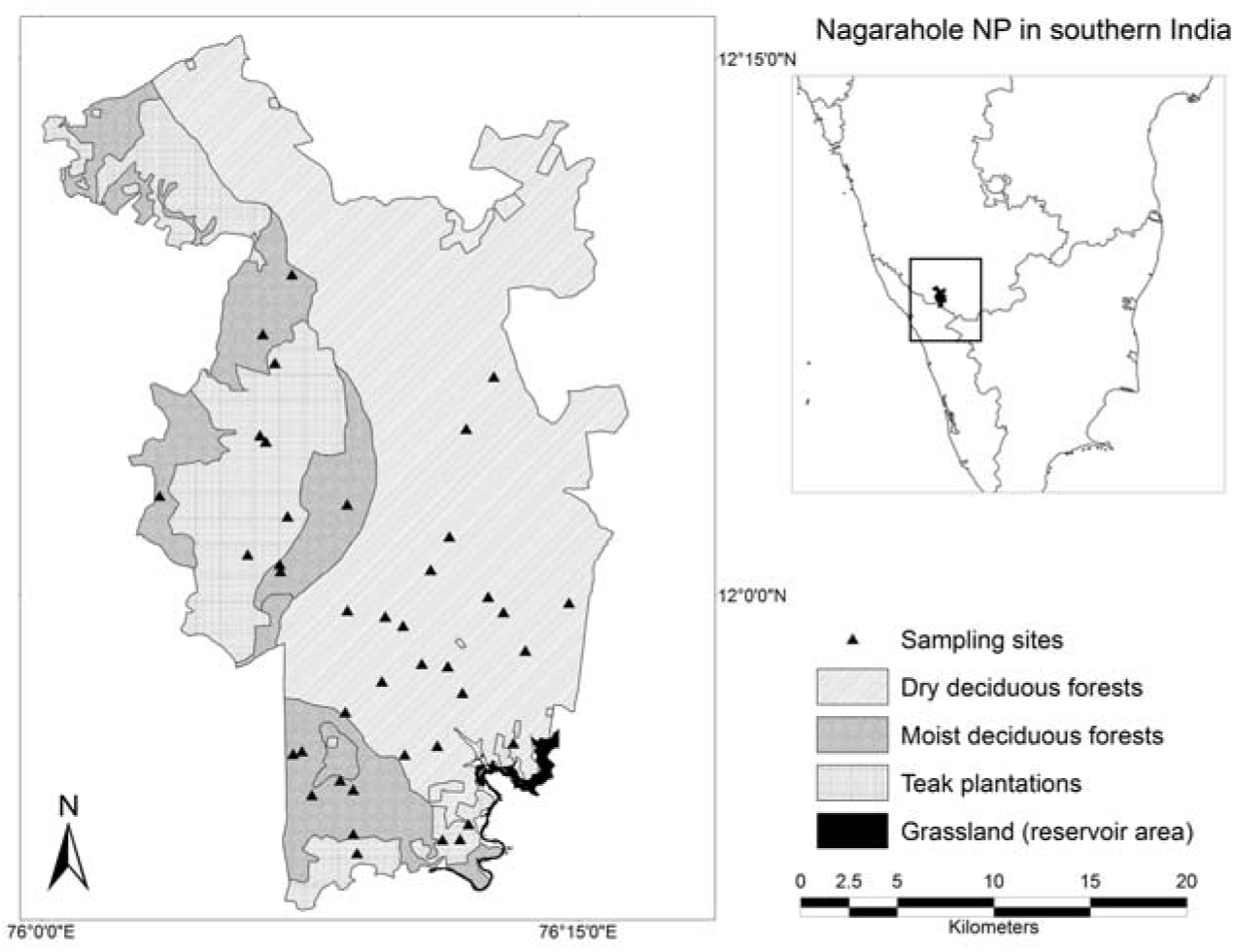
Locations of sampling sites in the study area. The forest type classification is based on Pascal (1982). Inset: map of southern India showing the location of Nagarahole National Park.

At the level of the graminoid community, two measures of visually-estimated overall graminoid abundance were used: total graminoid cover as described above, and the sum of species cover (the sum of individual graminoid species covers; the value might exceed 100% since each species was assessed independently; see Figure 1). Total graminoid biomass was normally distributed whereas individual species biomass data were non-normal and were, therefore, log-transformed for the analyses. However, the analyses were also performed on untransformed data to evaluate the robustness of the results. We first used homogeneity of regression slopes test (Zar 1974) to inspect the effect of forest type (dry deciduous, moist deciduous and teak forests) on the relationship between total graminoid biomass and overall graminoid cover. Similar relationships between total biomass and overall cover in different forest types would result in a homogeneity of slopes. We then performed multiple regressions of biomass on both estimates of overall graminoid cover to assess the utility of both measures in predicting total graminoid biomass. We also used the weighted average of graminoid species heights (weighted according to species cover) as an additional explanatory variable and performed multiple regressions to test the utility of height in improving total biomass estimates. Akaike information criterion with small sample correction (*AICc*; Hurvich and Tsai 1989) was used for selection from the regression models including and excluding height. Although there was a homogeneity of slopes (see Results) and plots from all forest types could be combined for further analyses, we used forest type as a factor to account for the off-chance that pooling the data would affect the results. In order to include the information on forest type in multiple regressions, two dummy categorical variables were generated: deciduous (category 1) or not (category 0 representing teak plantation), and moist deciduous (1) or not (0 representing dry deciduous forest when the previous categorical variable had value 1). These two variables were included in all regression analyses of data from the forest habitats. At the level of individual species, multiple regressions (including the variables for forest type) of species cover on species biomass were carried out for 10 common species (the other species were present in fewer than 10 plots) all of which happened to be elephant food species. We also checked whether multiple regressions that included the individual species’ height explained variation in individual species’ biomass better compared to regressions lacking this information. Model selection from the two types of regression models was done on the basis of *AICc*. All the regressions described above were carried out on data from two spatial scales: a) 1 m × 1 m quadrat level on which the measurements were originally made, and b) 20 m × 5 m plot level, such that the plots were spatially independent. Values of different variables in 1 m × 1 m quadrats were averaged to obtain values for the 20 m × 5 m plots.

We also carried out multiple regressions to find out how closely species diversity calculated using visually assessed cover data matched that calculated using biomass data. We calculated two commonly used measures of species diversity, Simpson’s diversity index and Shannon-Wiener index of diversity (see Southwood and Henderson 2009)). We calculated diversity of a) all the graminoid species and b) only elephant food species. Diversity index values from the three quadrats of every plot were averaged, and analyses were carried out using the average index value for each plot. Forest type variables were used as described above in all multiple regressions.

### Grassland data

Seven large stretches (called zones) across the length of the grassland were sampled. A zone would, therefore, be somewhat analogous to the 20 × 5 m plots in the forest, within which quadrats were sampled, although zones were much larger and quadrats within zones were randomly placed. Within each zone, 20 independent quadrats (1 m × 1 m) were marked, each chosen by walking from the centre of the zone up to a randomly chosen distance and along a randomly chosen direction (distance and angle obtained from a random number generator). Visual estimation of total graminoid cover was carried out in these quadrats and graminoid heights were measured before harvesting the above-ground biomass. Species-level data were not sampled since the only two grass species present could not be differentiated in their non-flowering states. Both these grasses are fed upon by elephants. The data were collected during four 30-day periods between mid-February and mid-June 2015 with equal numbers of random quadrats sampled in each period. Biomass measurements for 10 quadrats could not be performed because of heavy rain, which might have affected the weight significantly.

General regression models were used to analyse data from grassland quadrats. Total graminoid biomass was used as a dependent variable, total graminoid cover as an independent variable, and month and zone as categorical predictors to control for the effect of variation due to time and location, respectively. The analysis was done both with and without graminoid height as a continuous predictor.

Analyses were performed using Statistica 8 (StatSoft 2007).

## Results

### Relationship between visually-estimated total graminoid cover and total graminoid biomass in forest habitat

Based on the analysis of data from the 40 plots sampled in the three forest types, the mean graminoid biomass was 0.204 kg (95% CI: 0.165 to 0.241 kg). We found no effect of forest type on the relationship between total biomass and total cover, between total biomass and the sum of species covers, or between total biomass and weighted average of species heights (see Table 1). The statistics for regressions tests of how well the graminoid biomass is explained by cover and height are shown in Table 2. We found that both total graminoid cover and the sum of graminoid species cover were able to explain a large amount of variation in total biomass, with the sum of species cover showing a slightly higher coefficient of determination than the total cover (scatter-plots shown in Figure 3). Inclusion of weighted average of species heights to the multiple regression increased the coefficient of determination slightly in the case of sum of species cover and total cover compared to the respective models without the inclusion of height (Table 2). The small difference in *AICc* in both cases suggested that the models including height were not significantly better.

**Table 1:** Results from homogeneity of slopes test to examine the effect of forest type on the regression slopes of graminoid biomass on three continuous predictors: 1) sum of species covers, 2) total cover, and 3) weighted average of species heights.

|  | Sum of species<br>cover | Total cover | Weighted<br>heights |
| --- | --- | --- | --- |
| Effect of forest type<br>on slope | $F[2, 33] = 0.35,$<br>$P = 0.704$ | $F[2, 33] = 0.43,$<br>$P = 0.652$ | $F[2, 33] = 1.13,$<br>$P = 0.336$ |
| Effect of continuous<br>predictor | $F[1, 33] = 57.21,$<br>$P < 0.001$ | $F[1, 33] = 35.34,$<br>$P < 0.001$ | $F[1, 33] = 1.07,$<br>$P = 0.308$ |

**Table 2:**
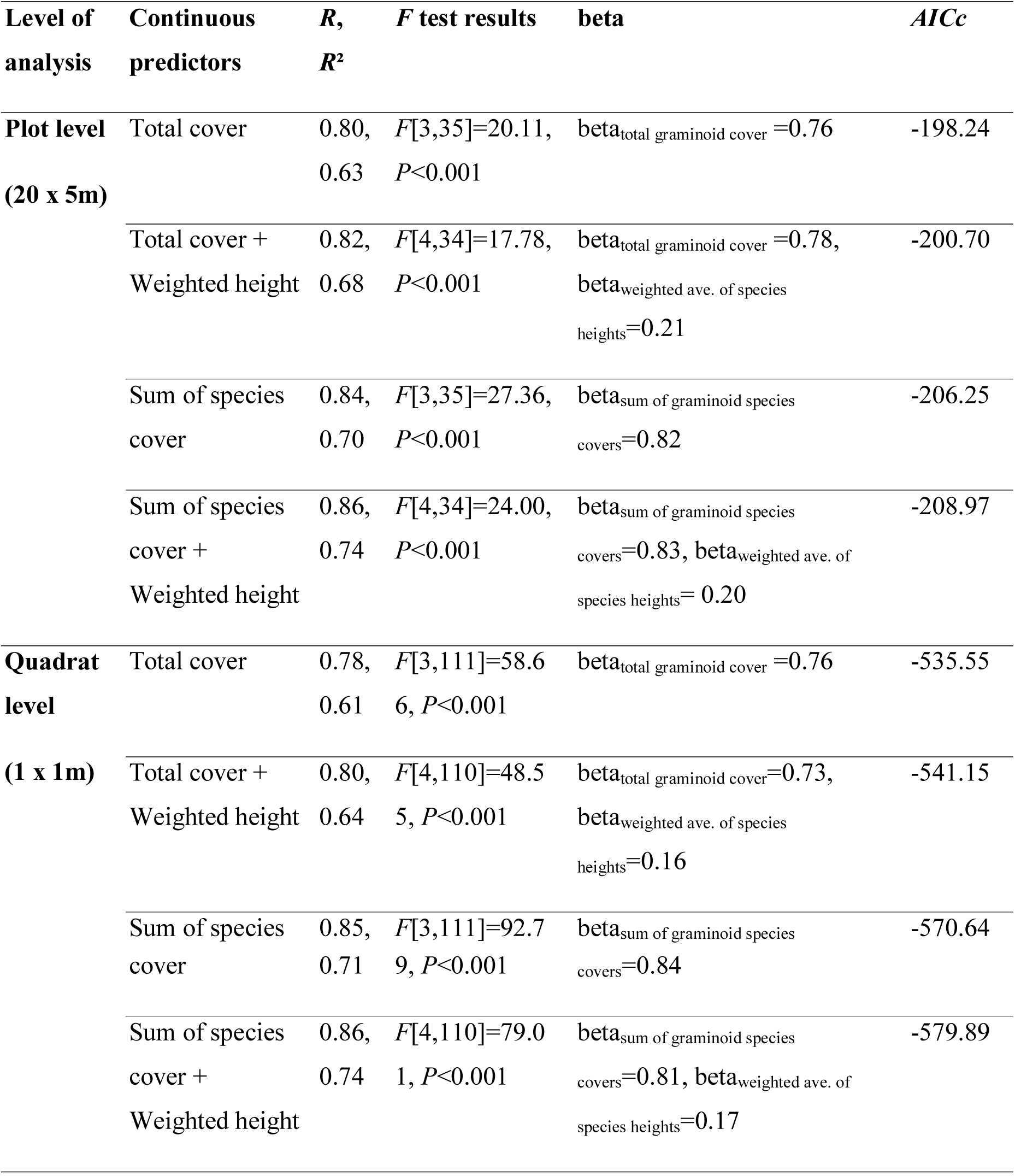
Results of regressions to examine how visual estimates of cover explain total graminoid biomass in forest habitat, and the additional utility of weighted average of species heights in improving the total biomass estimates. *P* < 0.05 for all beta coefficient values. *AICc* can be used to compare the regressions with and without average species height. The difference between a model with the lowest *AICc* and other models are not considered significant if the difference in *AICc* is less than two. The model with the lowest *AICc* is moderately better than the other models if the difference in *AICc* is between 4 and 7, and model with the lowest *AICc* is considerably better than the other models if the difference in *AICc* is >10.

**Figure 3:**
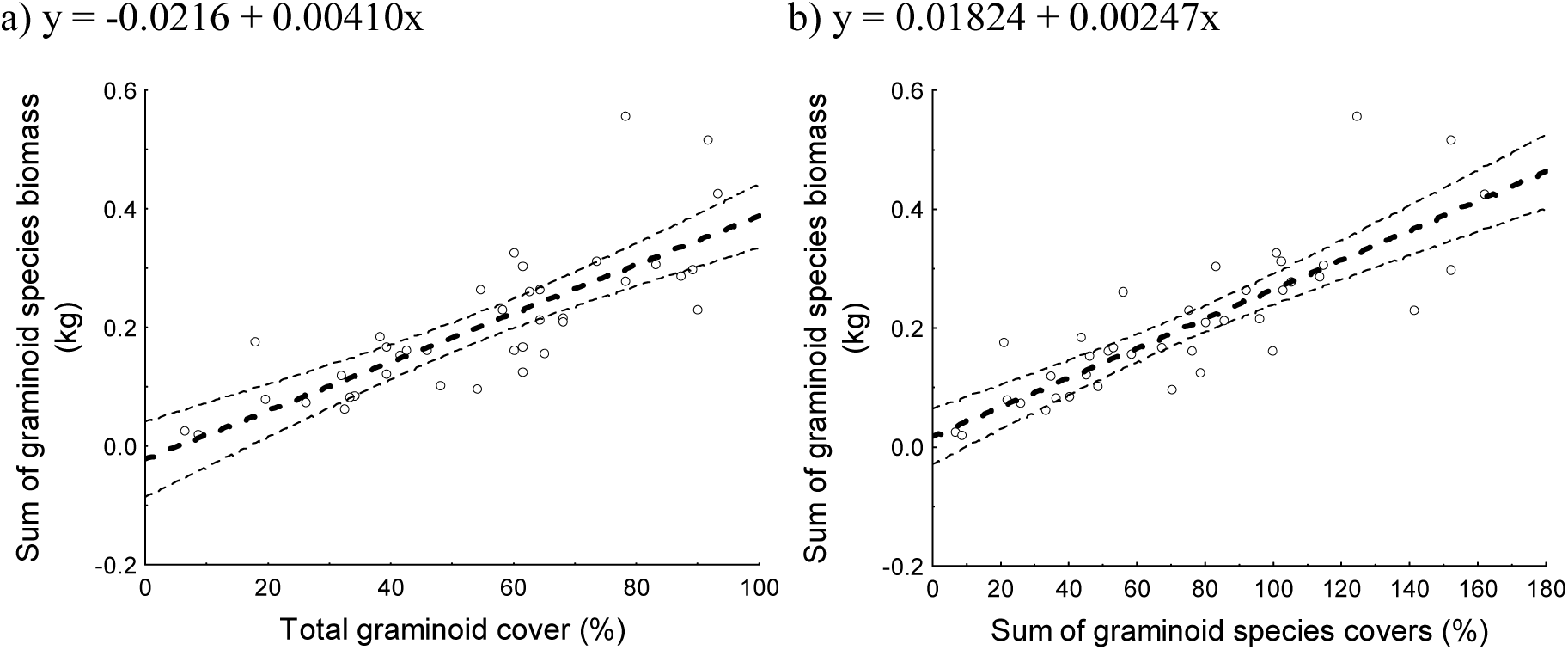
Scatter plots for the plot-level graminoid abundance data from the forest plots, showing the relation between a) visually-assessed total graminoid cover and total graminoid biomass and b) the sum of visually-assessed individual graminoid species cover and total graminoid biomass.

### Relationship between visually-estimated species covers of graminoid food species of elephants and their respective species biomass in forest habitat

Analyses of species-wise abundance at the plot level showed strong relationships between biomass and visually-estimated species cover for all the common (which were present in 10 or more plots) graminoid species (*R*^2^=0.63 to 0.98; Table 3). Multiple regressions using species average height as an additional predictor variable also yielded high *R*^2^ values (*R*^2^=0.69 to 0.98; Table 3), although average height had a significant effect in the regression only in the case of *Oplismenus compositus* and *Digitaria* sp.2 at the plot level (Table 3). With the exception of *Oplismenus compositus*, the *AICc* values for all other species tested were smaller when height was not included in the model to estimate biomass compared to the model when height was included along with cover. However, the differences in *AICc* values between the respective models were small, indicating that the models with and without heights largely performed equally well. At the quadrat level also (Table 4), high regression coefficients were obtained from the regression models that used species cover (*R*^2^=0.68 to 0.90) and the regressions that used both species cover and height (*R*^2^=0.68 to 0.90), with height having a significant effect in the case of *Cyrtococcum accrescens*, *Cyrtococcum oxyphyllum*, *Cyrtococcum patens*, *Oplismenus compositus*, and *Oryza sativa*. The *AICc* values were smaller in the models that included height in these species but none of the species showed large differences (>10) in AICc that would suggest an overwhelming advantage to adding height in regression models (Table 4).

**Table 3.** Results for plot-level regressions of graminoid species biomass on 1) visually-estimated species cover, and 2) visually-estimated species cover and measured average species height, in forest habitat. Details of the regressions are shown based on analysis of log-transformed data along with *R*^2^ untransformed data (shown as *R*^2^ (untr.)). *P* values for the regressions are not shown separately, except for *Digitaria* sp.2, because all the other *P* values (for transformed and untransformed data) were smaller than 0.001. #Effect of habitat in multiple regression was significant. **Effect of height was significant. See Table 2 for interpretation of *AICc* differences.

| Species | Species cover |  |  |  |  |  | Species cover and avg. species height |  |  |  |  |  |
| --- | --- | --- | --- | --- | --- | --- | --- | --- | --- | --- | --- | --- |
| | $F$ | $df_1, df_2$ | $R$ | $R^2$ | $AICc$ | $R^2$ (untr.) | $F$ | $df_1, df_2$ | $R$ | $R^2$ | $AICc$ | $R^2$ (untr.) |
| *Axonopus compressus* | 22.36 | 3,15 | 0.90 | 0.82 # | -57.69 | 0.71 | 15.77 | 4,14 | 0.90 | 0.82 # | -54.56 | 0.71 |
| *Chloris dolichostachya* | 73.62 | 3,14 | 0.97 | 0.94 # | -58.16 | 0.93 # | 56.32 | 4,13 | 0.97 | 0.95 # | -56.40 | 0.94 # |
| *Cynodon dactylon* | 58.58 | 3,10 | 0.97 | 0.95 | -42.22 | 0.83 | 39.93 | 4,9 | 0.97 | 0.95 | -38.30 | 0.85 |
| *Cyrtococcum accrescens* | 38.47 | 3,22 | 0.92 | 0.84 | -65.74 | 0.67 # | 29.80 | 4,21 | 0.92 | 0.85 | -64.67 | 0.69 |
| *Cyrtococcum oxyphyllum* | 56.99 | 3,16 | 0.96 | 0.91 | -58.39 | 0.87 | 41.34 | 4,15 | 0.96 | 0.92 | -55.79 | 0.88 |
| *Cyrtococcum patens* | 113.81 | 3,8 | 0.99 | 0.98 | -40.45 | 0.89 | 82.59 | 4,7 | 0.99 | 0.98 | -36.97 | 0.89 |
| *Digitaria* sp.2 | 4.03 | 3,7;$P=0.059$ | 0.80 | 0.63 | -23.69 | 0.74 | 3.33 | 4,6;$P=0.092$ | 0.83 | 0.69 | -20.29 | 0.90** |
| *Kyllinga monocephala* | 82.01 | 3,15 | 0.97 | 0.94 | -48.02 | 0.89 | 57.55 | 4,14 | 0.97 | 0.94 | -44.81 | 0.89 |
| *Oplismenus compositus* | 74.09 | 3,33 | 0.93 | 0.87 | -100.10 | 0.79 | 64.40 | 4,32 | 0.94 | 0.89** | -103.38 | 0.84** |
| *Oryza sativa* | 21.02 | 3,10 | 0.93 | 0.86 | -36.39 | 0.86 | 16.17 | 4,9 | 0.94 | 0.88 | -33.94 | 0.87 |

**Table 4.** Results for quadrat-level regressions of graminoid species biomass on visually-estimated species cover, and of species biomass on visually-estimated species cover and measured average species height for forest habitat. Details of the regressions are shown based on analysis of log-transformed data along with *R*^2^ untransformed data (shown as *R*^2^ (untr.)). P<0.001 for multiple regressions. #Effect of habitat in multiple regression was significant. **Effect of height was significant. See Table 2 for interpretation of *AICc* differences.

| Species | Species cover |  |  |  |  |  | Species cover and avg. species height |  |  |  |  |  |
| --- | --- | --- | --- | --- | --- | --- | --- | --- | --- | --- | --- | --- |
| | $F$ | $df_1, df_2$ | $R$ | $R^2$ | $AICc$ | $R^2$ (untr.) | $F$ | $df_1, df_2$ | $R$ | $R^2$ | $AICc$ | $R^2$ (untr.) |
| *Axonopus compressus* | 25.84 | 3,26 | 0.87 | 0.75 | -61.57 | 0.76 | 19.79 | 4,25 | 0.87 | 0.76 | -60.24 | 0.76 |
| *Chloris dolichostachya* | 36.68 | 3,32 | 0.88 | 0.77 | -83.80 | 0.88 | 27.18 | 4,31 | 0.88 | 0.78 | -81.81 | 0.88 |
| *Cynodon dactylon* | 55.33 | 3,21 | 0.94 | 0.89 | -67.24 | 0.83 | 45.49 | 4,20 | 0.95 | 0.90 | -67.53 | 0.83 |
| *Cyrtococcum accrescens* | 50.27 | 3,41 | 0.89 | 0.79 # | -98.43 | 0.53 | 39.72 | 4,40 | 0.89 | 0.80 | -98.76 | 0.61** |
| *Cyrtococcum oxyphyllum* | 59.39 | 3,38 | 0.91 | 0.82 | -100.92 | 0.70 | 52.63 | 4,37 | 0.92 | 0.85** | -105.28 | 0.72 |
| *Cyrtococcum patens* | 42.47 | 3,16 | 0.94 | 0.89 | -55.75 | 0.74 | 31.50 | 4,15 | 0.95 | 0.89 | -53.53 | 0.81** |
| *Digitaria* sp.2 | 10.69 | 3,15 | 0.83 | 0.68 | -26.19 | 0.81 | 7.48 | 4,14 | 0.83 | 0.68 | -22.94 | 0.81 |
| *Kyllinga monocephala* | 81.55 | 3,28 | 0.95 | 0.90 | -78.76 | 0.81 | 63.29 | 4,27 | 0.95 | 0.90 | -78.17 | 0.82 |
| *Oplismenus compositus* | 91.64 | 3,82 | 0.88 | 0.77 | -170.46 | 0.73 | 73.76 | 4,81 | 0.89 | 0.78** | -173.8 | 0.74** |
| *Oryza sativa* | 24.31 | 3,14 | 0.92 | 0.84 | -43.74 | 0.85 | 27.39 | 4,13 | 0.94 | 0.89** | -47.90 | 0.89** |

### Relationship between diversity indices calculated from visually-estimated species covers and the respective species biomass in forest habitat

The average number of graminoid species per 1 m × 1 m quadrat was 4.7 (95% CI: 4.02–5.38) out of which the average number of food species per quadrat was 3.74 (95% CI: 3.24–4.24). Species diversity calculated using visually-estimated cover explained a large proportion of variance in the diversity calculated using biomass data, at both the plot level (20 m × 5 m) and quadrat level, and when diversity was measured by either the Simpson’s diversity index or the Shannon-Wiener diversity index (*H′*) (Figure 4, Table 5). Similar regressions of biomass-based diversity on visual cover-based diversity using only elephant food graminoid species, rather than all graminoid species, also showed strong relationships (Table 5).

**Figure 4:**
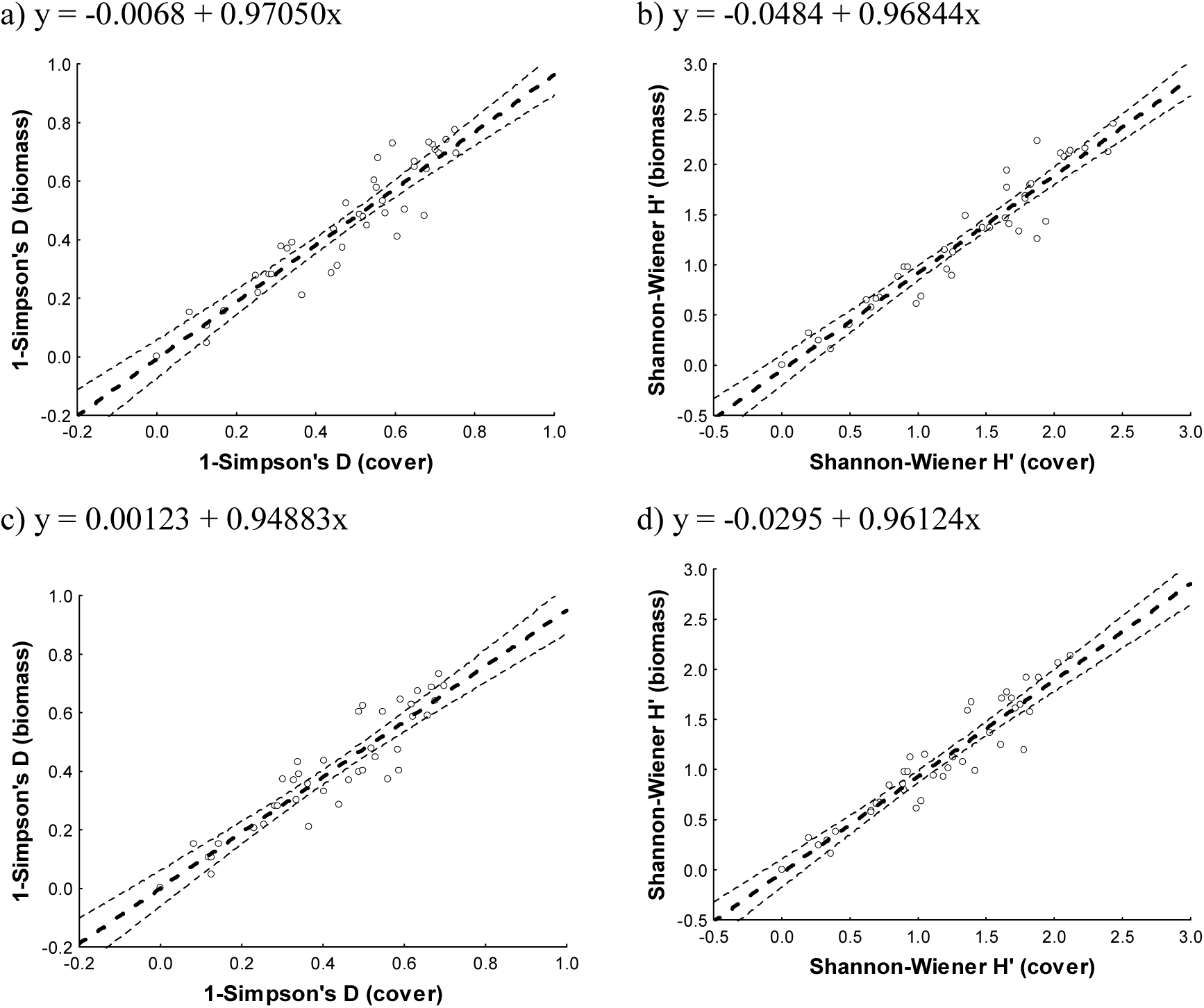
Scatter plots for the plot-level graminoid abundance data from the forest plots, showing the relationships between Simpson’s diversity indices (1-*D*) calculated from cover and biomass data (a, c) and Shannon-Wiener diversity indices (*H’*) calculated from cover and biomass data (b, d). All graminoid species are included in a) and b), while only elephant food species are included in c) and d).

**Table 5:** Results from multiple regressions to examine the relationship between diversity indices calculated using biomass data and cover data. *P* < 0.001 in all cases.

| Scale | All graminoid species |  | Food graminoid species |  |
| --- | --- | --- | --- | --- |
|  | Simpson's index | Shannon-Wiener | Simpson's index | Shannon-Wiener |
|  | (1-D) | index (H') | (1-D) | index (H') |
| *Plot Level* | $R=0.93, R^2=0.87,$<br>$F[3,36]=79.92$ | $R=0.85, R^2=0.72,$<br>$F[3,36]=30.12$ | $R=0.92, R^2=0.85,$<br>$F[3,36]=67.42$ | $R=0.95, R^2=0.89,$<br>$F[3,36]=101.44$ |
| *Quadrat Level* | $R=0.89, R^2=0.80,$<br>$F[3,115]=150.47$ | $R=0.93, R^2=0.86,$<br>$F[3,115]=232.56$ | $R=0.88, R^2=0.78,$<br>$F[3,115]=132.86$ | $R=0.92, R^2=0.84,$<br>$F(3,115)=205.55$ |

### Relationship between visually-estimated total graminoid cover and total graminoid biomass in grassland habitat

Based on data from 550 quadrats, the mean graminoid biomass was calculated to be 0.684 kg (95% CI: 0.655–0.712 kg). Visual estimation of total graminoid cover, along with month and zone ID as categorical predictors, explained total graminoid biomass to a large extent (General regression model, test of SS Whole Model vs. SS Residual: *R* = 0.78, *R^2^*= 0.61, *F*[3, 54] = 281.64, *P* < 0.001, *AICc* = -1687.49). The effects of total graminoid cover and month were significant (beta_total graminoid cover_ = 0.51, beta_month_ = 0.41), whereas zone did not have a significant effect. The addition of average height improved the *R*^2^ value (Test of SS Whole Model vs. SS Residual: *R* = 0.83, *R^2^* = 0.69, *F*[4, 544] = 306.84, *P* < 0.001, *AICc* = -1819.85), with significant effects of total graminoid cover, height, and month, but not zone, on biomass (beta_total graminoid cover_ = 0.44, beta_height_ = 0.30, beta_month_ = 0.41).

## Discussion

We found that visual assessment of cover, which allows for rapid sampling, performed very well in assessing forage availability in forest and grassland habitat. Using this method, we were able to obtain fairly accurate estimates of biomass of graminoids in general and food graminoids of Asian elephants in particular. The biomass harvest method has been suggested to be an ideal measure of abundance (Wilson 1991, Chiarucci et al. 1999) of herbaceous vegetation, but can be time-consuming if species have to be hand-sorted and weighed separately (Harmony et al. 1997, Lavorel et al. 2008). Biomass harvesting also has limitations with respect to permits in critical wildlife areas because of its destructive nature, unlike visual estimation of cover. We found that visually estimated cover, which does not suffer from these drawbacks, was a good proxy for biomass of individual food species, as well as total graminoid biomass, in complex forest habitats. Such estimation of individual species or portions of the vegetation (in our study area, herbs are more abundant than graminoids) is not feasible with other non-destructive methods (discussed in Radloff and Mucina 2007, Redjadj et al. 2012, Walter et al. 2015). An objection to the visual estimation of species covers had been that the “sum of species covers” as a measure of total cover would give “non-sense number”s that exceeded 100% (Wilson 2011). Wilson (2011) had argued that there was no particular reason why between-species leaf overlaps would be helpful while within-species overlaps would not be important. However, empirically, the sum of species covers performed better at explaining total graminoid biomass in our study, and, perhaps, this is because it incorporates at least some (between-species) overlap component (see Figure 1), unlike the total graminoid cover that does not include any overlap. The sum of graminoid species covers may perform better than total graminoid cover when the within-species leaf overlap is smaller than the between-species leaf overlap. We speculate that this might be true of forests with multiple strata, in which individuals in the lower strata avoid self-shading and individuals of the same species are not very close to one another in order to reduce competition. Since the sum of graminoid species cover better represents total biomass compared to total cover (as discussed above) and since species-level cover estimates are highly related to species biomass, one can obtain good estimates of the proportional abundance of foods by dividing the sum of food species covers by the sum of all species covers. This would be useful for studies on foraging ecology in tropical forests with high diversity where food species are a small fraction of the total number of species present.

The inclusion of height in regression models yielded mixed results. In the forests, total graminoid biomass and sum of species biomass were explained adequately by cover and most models that included height were not substantially better than those that did not include height (small differences in *AICc*). The effect of height was not significant at the individual species-level also for most food species. On the other hand, the relationship between cover and total graminoid biomass was improved by the inclusion of average height in the grassland habitat. This pattern may result from higher variability in total cover (average total graminoid cover=54.85%, CV total graminoid cover=42.33%) than in height (average weighted height=24.63cm, CV weighted height=32%) in forest habitat. As cover by graminoids saturates (total cover estimate has an upper limit of 100% whereas biomass and height are not limited) and relative variability in cover decreases vis-à-vis height, for example, in grasslands or swamps, the explanatory power of height with respect to that of cover is expected to increase. Thus, in the grassland habitat, where cover is closer to saturation and less variable (average total graminoid cover: 87.27%, CV for total graminoid cover: 19.17%; average height=5.71cm, CV for height: 46.84%) the effect of height relative to that of graminoid cover (beta_total graminoid cover_=0.44, beta_height_ =0.30, *P*<0.05 for both) was greater than that in the forest habitat (beta_total graminoid cover_=0.78, beta_weighted average of species heights_=0.21, *P*<0.05 for both). We thus found that there was no overwhelming benefit to including height in forest habitat, while it might be worthwhile measuring height in grassland habitat. The improvement in biomass estimation models after inclusion of height has also been reported in other grassland habitats like tussock-grasslands in alpine Andes (Oliveras et al. 2013) and rangelands in Argentina (Guevara et al. 2002). In forest habitats, such relationships have been rarely studied for graminoids, although in one study in pine forests of Arizona, height did not result in substantial improvement to predictive power of models that used only cover (Andariese and Covington 1986). In habitats where cover is near saturation value (100%), we suggest that measurement of height can substantially improve biomass estimates, as seen in grassland habitat in our study. The limited predictive power of cover in plots with high cover values (above 80%) has also been discussed in Axmanová et al. (2012) for wet meadows of temperate Central Europe. One caveat is that we do not know if graminoid height itself affects selection of foraging sites by animals.

We also found that the species diversity of graminoids as calculated from cover data explained large variations in the diversity indices measured from biomass data when used along with forest type as a categorical variable. This similarity in community characteristics from the two methods further support the utility of visual estimation as a rapid assessment method in foraging ecology since ecologists may also be interested in assessing the diversity of foraging sites in order to test whether an animal selects a few food items or feeds on a wide range of species from the available plant species (e.g. Owen-Smith and Chafota 2012). Apart from selectivity, since prolonged feeding on a few plant species may result in accumulation of specific secondary metabolites which can be difficult to detoxify by the mechanism for those metabolites, it is hypothesised that feeding on a wide range of species allows multiple counter pathways to share the load of detoxification of multiple types of secondary metabolites (reviewed in Marsh et al. 2006). Some generalist feeders may thus adopt a strategy of feeding on a variety of plants, thereby increasing their feeding rates. Quantification of species diversity may thus be useful in addressing such proximal aspects of foraging which affect patch selection. Such proximate mechanisms have been invoked to explain the relation between the diet of African elephants and tree diversity in the habitat (Cordon et al. 2011).

One possible limitation of this study is that we measured fresh rather than dry biomass. Fresh biomass has a component of leaf water content, which can change temporally based on environmental conditions. Therefore, the cover-biomass relationship that we recovered based on fresh biomass measurement may not be the same as that based on dry biomass measurement. However, it is often of interest to measure fresh biomass in studies of foraging because that is the weight of food that an animal would consume. We addressed the issue of observer bias associated with the visual method (Tonteri 1990) to some extent by having a single observer (HG) carry out the cover estimation from all sites. As observer-related errors can cumulatively become large if many observers are involved in data collection (Tonteri 1990, Klimeš 2003), it would be necessary to consider the error across observers before the final estimation.

In summary, we find that the visual estimation method performs very well in assessing forage availability in a tropical forest and a grassland habitat, and can, therefore, be used in studies of elephant habitat and forage selection. This will save time and allow for sampling a larger number of sites. Our study was carried out in tropical deciduous forests, which constitute more than 65% of the total forest area in India (Reddy et al. 2015) and about one-sixth of the forest cover in South-east Asia (Wohlfart et al. 2014). It is likely that the positive relationship between cover and biomass will hold in similar forests, although the strength of the relationship may vary geographically. We do not imply that the relationships we find are completely transferable to other locations and suggest independent assessments in order to develop site-specific cover-to-biomass models. However, since we show that visual estimates of cover can be very useful in studies of foraging, this opens up the method for use by various researchers, who may have otherwise been deterred from using this based on a few previous studies. Moreover, since other sympatric ungulates like *Axis axis* and *Bos gaurus* are also primarily grazers (Ahrestani et al. 2012), the visual estimation method should also work well for quantifying the resource distribution for these generalist ungulates in our study site and similar deciduous forests.

## Acknowledgements

This work was funded by the Department of Science and Technology’s (Government of India) Ramanujan Fellowship to TNCV, the Council of Scientific and Industrial Research, Government of India, and National Geographic Society, USA. JNCASR provided logistic support. We thank the office of the PCCF, Karnataka Forest Department, for field permits, and various officials and staff of Nagarahole National Park for permits and support at the field site. Ranga, Shankar and Krishna provided field assistance. We thank two anonymous reviewers and the handling editor for their comments and suggestions.

## References

Ahrestani, F.S., Heitkönig, I.M. and Prins, H.H. 2012. Diet and habitat–niche relationships within an assemblage of large herbivores in a seasonal tropical forest. Journal of Tropical Ecology 28: 385–394.

Andariese, S.W. and Covington, W.W. 1986. Biomass estimation for four common grass species in northern Arizona ponderosa pine. Journal of Range Management 39: 472–473.

Axmanová, I., Tichý, L., Fajmonová, Z., Hájková, P., Hettenbergerová, E., Li, C.F., Merunková, K., Nejezchlebová, M., Otýpková, Z., Vymazalová, M. and Zelený, D. 2012. Estimation of herbaceous biomass from species composition and cover. Applied Vegetation Science 15: 580–589.

Baskaran, N., Balasubramanian, M., Swaminathan, S. and Desai, A.A. 2010. Feeding ecology of the Asian elephant *Elephas maximus* Linnaeus in the Nilgiri Biosphere Reserve, southern India. Journal of the Bombay Natural History Society 107: 3–13.

Blake, S., 2003. The Ecology of Forest Elephant Distribution and its Implications for Conservation. PhD thesis, University of Edinburgh, Edinburgh, UK.

Chiarucci, A.J.B.W., Wilson, J.B., Anderson, B.J. and Dominicis, V. 1999. Cover versus biomass as an estimate of species abundance: does it make a difference to the conclusions? Journal of Vegetation Science 10: 35–42.

Codron, D., Lee-Thorp, J.A., Sponheimer, M., Codron, J., De Ruiter, D. and Brink, J.S. 2007. Significance of diet type and diet quality for ecological diversity of African ungulates. Journal of Animal Ecology 76: 526–537.

Drew, W.B., 1944. Studies on the use of the point-quadrat method of botanical analysis of mixed pasture vegetation. Journal of Agricultural Research 69: 289–297.

Elzinga, C.L., Salzer, D.W. and Willoughby, J.W. 1998. Measuring and Monitoring Plant Populations. Bureau of Land Management, Denver, Colorado.

Guevara, J.C., Gonnet, J.M. and Estevez, O.R. 2002. Biomass estimation for native perennial grasses in the plain of Mendoza, Argentina. Journal of Arid Environments 50: 613–619.

Guo, Q. and Rundel, P.W. 1997. Measuring dominance and diversity in ecological communities: choosing the right variables. Journal of Vegetation Science 8: 405–408.

Harmoney, K.R., Moore, K.J., George, J.R., Brummer, E.C. and Russell, J.R. 1997. Determination of pasture biomass using four indirect methods. Agronomy Journal 89: 665–672.

Henschel, J.R., Burke, A. and Seely, M. 2005. Temporal and spatial variability of grass productivity in the central Namib Desert. African Study Monographs 30: 43–56.

Hermy, M., 1988. Accuracy of visual cover assessments in predieting standing crop and environmental correlation in deciduous forests. Vegetatio 75: 57–64.

Hurvich, C.M. and Tsai, C.L. 1989. Regression and time series model selection in small samples. Biometrika 76: 297–307.

Hutto, R. L. 1990. Measuring the availability of resources. Studies in Avian Biology 13: 20–28.

Iversen, M., Fauchald, P., Langeland, K., Ims, R. A., Yoccoz, N. G. and Bråthen, K. A. 2014 Phenology and cover of plant growth forms predict herbivore habitat selection in a high latitude ecosystem. PLoS One 9: 1–11.

Kennedy, K.A. and Addison, P.A. 1987. Some considerations for the use of visual estimates of plant cover in biomonitoring. Journal of Ecology 75: 151–157.

Klimeš, L. 2003. Scale-dependent variation in visual estimates of grassland plant cover. Journal of Vegetation Science 14: 815–821.

Lavorel, S., Grigulis, K., McIntyre, S., Williams, N. S. G., Garden, D., Dorrough, J., Berman, S., Quétier, F., Thébault, A. and Bonis, A. 2008. Assessing functional diversity in the field - methodology matters! Functional Ecology 22: 134–147.

Marsh, K.J., Wallis, I.R., Andrew, R.L. and Foley, W.J. 2006. The detoxification limitation hypothesis: where did it come from and where is it going? Journal of Chemical Ecology 32: 1247–1266.

Noyce, K. V. and Coy, P. L. 1990. Abundance and productivity of bear food species in different forest types of northcentral Minnesota. In Bears: Their Biology and Management, Vol. 8, A Selection of Papers from the Eighth International Conference on Bear Research and Management, Victoria, British Columbia, 1990, pp. 169–181.

Oliveras, I., Eynden, M., Malhi, Y., Cahuana, N., Menor, C., Zamora, F. and Haugaasen, T. 2014. Grass allometry and estimation of above-ground biomass in tropical alpine tussock grasslands. Austral Ecology 39: 408–415.

Owen-Smith, N. and Chafota, J. 2012. Selective feeding by a megaherbivore, the African elephant (*Loxodonta africana*). Journal of Mammalogy 93: 698–705.

Owen-Smith, R. N. 1988. Megaherbivores: The Influence of Very Large Body Size on Ecology. Cambridge University Press, Cambridge, UK.

Pascal, J. P. 1982. Bioclimates of the Western Ghats: Maps 1-2. French Institute of Pondicherry, Pondicherry, India.

Pekin, B.K., Endress, B.A., Wisdom, M.J., Naylor, B.J. and Parks, C.G. 2015. Impact of ungulate exclusion on understorey succession in relation to forest management in the Intermountain Western United States. Applied Vegetation Science 18: 252–260.

Pettorelli, N., Ryan, S.J., Mueller, T., Bunnefeld, N., Jedrzejewsk, B., Lima, M. and Kausrud, K., 2011. The Normalized Difference Vegetation Index (NDVI): unforeseen successes in animal ecology. Climate Research 46: 15–27.

Radloff, F.G.T. and Mucina, L. 2007. A quick and robust method for biomass estimation in structurally diverse vegetation. Journal of Vegetation Science 18: 719–724.

Rebollo, S., Milchinas, D. G., Stapp, P., Augustine, D. J. and Derner, J. D. 2013. Disproportionate effects of non-colonial small herbivores on structure and diversity of grassland dominated by large herbivores. Oikos 122: 1757–1767.

Reddy, C.S., Jha, C.S., Diwakar, P.G. and Dadhwal, V.K. 2015. Nationwide classification of forest types of India using remote sensing and GIS. Environmental Monitoring and Assessment 187: 1–30.

Redjadj, C., Duparc, A., Lavorel, S., Grigulis, K., Bonenfant, C., Maillard, D., Saïd, S. and Loison, A. 2012. Estimating herbaceous plant biomass in mountain grasslands: a comparative study using three different methods. Alpine Botany 122: 57–63.

Saïd, S., Pellerin, M., Guillon, N., Débias, F. and Fritz, H. 2005. Assessment of forage availability in ecological studies. European Journal of Wildlife Research 51: 242–247.

Southwood, T.R.E. and Henderson, P.A., 2009. Ecological methods. John Wiley & Sons.

StatSoft, Inc. 2007. Statistica (data analysis software system), Version 8.0. www.statsoft.com

Sukumar, R. 1990. Ecology of the Asian elephant in southern India. II. Feeding habits and crop raiding patterns. Journal of Tropical Ecology 6: 33–53.

Tonteri, T. 1990. Inter-observer variation in forest vegetation cover assessments. Silva Fennica 24: 189–196.

Vidya, T.N.C., Prasad, D. and Ghosh, A. 2014. Individual identification in Asian elephants. Gajah 40: 3–17.

Walter, C.A., Burnham, M.B., Gilliam, F.S. and Peterjohn, W.T. 2015. A reference-based approach for estimating leaf area and cover in the forest herbaceous layer. Environmental Monitoring and Assessment 187: 1–9.

Wilmshurst, J.F., Fryxell, J.M. and Colucci, P.E. 1999. What constrains daily intake in Thomson’s gazelles? Ecology 80: 2338–2347.

Wilson, J.B. 1991. Methods for fitting dominance/diversity curves. Journal of Vegetation Science 2: 35–46.

Wilson, J.B. 2011. Cover plus: ways of measuring plant canopies and the terms used for them. Journal of Vegetation Science 22: 197–206.

Wohlfart, C., Wegmann, M. and Leimgruber, P. 2014. Mapping threatened dry deciduous dipterocarp forest in South-east Asia for conservation management. Tropical Conservation Science 7: 597–613.

Zar, J.H. 1974. Biostatistical Analysis. Prentice-Hall, NJ.

